# Pangenome-guided interpretation of *k*-mer GWAS in *Capsicum*

**DOI:** 10.64898/2026.11.18.764533

**Authors:** Christina Papastolopoulou, Patrick Flege, Peter Polacek, Yury Tikunov, Arnaud Bovy, Dick de Ridder, Sander A. Peters, Sandra Smit

**Affiliations:** Wageningen Plant Research, Bioscience, cluster Applied Bioinformatics, Wageningen Research, Droevendaalsesteeg 1, Wageningen, 6708 PB, The Netherlands; Bioinformatics Group, Wageningen University & Research, Droevendaalsesteeg 1, Wageningen, 6708 PB, The Netherlands; Breeding for Food Quality, Wageningen Plant Research, Droevendaalsesteeg 1, Wageningen, 6708 PB, The Netherlands

**Keywords:** GWAS, *k*-mer, pangenome, *Capsicum*

## Abstract

Agronomic trait diversity in *Capsicum* is shaped by sequence variation that it is not always captured by SNPs called against a single reference genome. Reference-free *k*-mer GWAS can detect variants beyond SNPs, but significant *k*-mers are difficult to interpret as they lack genomic coordinates and annotation. In this study, we developed a pangenome-guided workflow to localize significant *k*-mers in a graph-based *Capsicum* pangenome. The workflow connects significant *k*-mers, identified through *k* - mer GWAS, to genomic positions, their annotations and homology groups. We applied this approach to a metabolite dataset of 343 *C. annuum* accessions, focusing on the cases of capsaicinoids and specific aroma compounds. For capsaicinoid content, our workflow provided insight in the haplotype composition of the sequenced accessions in the *Pun1* locus and the genomic variations contributing to pungency. In the case of dehydroxylinalool oxide, the analysis identified that the haplotype associated to high phenotypes contained an extra terpene synthase (*TPS*). And last, the production of ***α***-ylangene seems to be influenced by a 2 Mb inversion on chromosome 9, right next to a terpene synthase gene. Integrating *k*-mer GWAS with graph-pangenome localization improves the interpretation of *k*-mer association signals in complex plant genomes. This workflow can help identify structural variation, haplotypes and candidate loci relevant for trait/metabolic diversity and future breeding applications.

## 1 Introduction

Population genomics has evolved into a practical approach for exploring the genetic origin of complex traits in both model organisms and crops [1]. High-throughput sequencing and scalable algorithms now enable association analyses across many genomes, revealing a wide range of polymorphisms underlying phenotypic diversity, including single-nucleotide variants, structural variants, presence-absence variation (PAV), and accessory gene content [2]. The shift from single-reference genomes to population-level genome representations is particularly important for crops such as *Capsicum*, where domestication, breeding, and transposable element activity have generated extensive structural and gene-content variation that a single linear genome reference cannot entirely capture [3–5].

To exploit the genomic diversity in *Capsicum* for trait discovery, reliable statistical frameworks are required to associate genetic variation with phenotypic outcomes. The genome-wide association study (GWAS) remains one of the primary statistical approaches to link genotypic to phenotypic variation [1, 6]. Conventional GWAS based on single-nucleotide polymorphisms (SNPs), however, is inherently reference-based, only considering variations relative to the chosen reference. This can limit the detection of biologically relevant variation, including large structural variants, presence-absence variation, and rearrangements that are absent from the reference [7, 8]. Reference-free *k*-mer GWAS approaches address part of this reference bias by associating short sequences (*k*-mers) directly with traits, bypassing variant calling and enabling the detection of multiple types of sequence variation from sequencing reads [7]. Scalable *k*-mer association methods were first widely applied in microbial genomics, where they helped identify trait-associated variation in highly variable genomes, and have since been adopted in plant studies to capture sequence variation that is difficult to represent through SNP-based approaches alone [9–13].

Despite its power, *k*-mer GWAS output is difficult to interpret, because a significant *k*-mer is only a short sequence that lacks genomic coordinates, annotation, and haplotype context [7, 12]. Aligning the significant *k*-mers to a single reference genome will reintroduce reference bias, particularly when the associated sequences originate from alternative alleles or structural variants that are absent from the selected reference. Alternatively, *de novo* assembly of overlapping significant *k*-mers, or the reads containing them, can be unreliable in repetitive or structurally complex regions, where assemblies may be fragmented or ambiguously localized. Thus, while *k*-mer GWAS removes one layer of bias from discovery, it creates a need for localization and interpretation strategies that can cluster associated *k*-mers into relevant/coherent loci and place them in a genomic context suitable for biological interpretation and breeding analyses.

Graph pangenomes offer a way to add genomic context to reference-free *k*-mer association signals. Existing *k*-mer GWAS methods differ in how *k*-mer variation is measured, ranging from binary presence-absence patterns across accessions to abundance models [7, 14–16]. By representing multiple genomes in a single non-linear structure, graph pangenomes capture sequence diversity, structural variation, and gene-content differences that are not fully represented by a single reference genome. Their adoption in plant genomics has grown as pangenome projects have revealed extensive gene-content diversity and structural rearrangements in crops such as *Capsicum*, highlighting the need for genome representations that better reflect population-level variation [4, 5, 17, 18]. Recent plant *k*-mer GWAS studies have started to reduce reference bias by aligning significant *k*-mers to multiple genomes or pangenome resources, demonstrating the value of interpreting association signals beyond a single reference [12, 13]. Graph-based approaches such as NodeGWAS perform association testing directly on pangenome graph nodes, improving inter-pretability but requiring a representative graph pangenome and reliable graph-node genotyping [19]. However, methods that maintain reference-free *k*-mer GWAS as the discovery step while linking significant *k*-mers to genome-specific coordinates, annotations, homology, and syntenic context in complex plant pangenomes remain limited.

Here, we use reference-free *k*-mer GWAS to identify sequence variation associated with secondary metabolite production in *Capsicum annuum*. To interpret these association signals, we present a workflow that localizes significant *k*-mers directly in a graph-based pangenome and connects them to interpretable genomic regions. Our approach integrates *k*-mer association analysis [7] with the PanTools pangenome framework [20, 21] to identify, cluster, and interpret trait-associated loci represented by enriched *k*-mer signals. By avoiding dependency on a single linear reference, this workflow enables association signals to be interpreted at the level of genome-specific loci, annotations, syntenic regions, and haplotypes across the genetic diversity of the population. We apply the pipeline to a graph pangenome representing major *Capsicum* lineages and accessions, and show how graph-localized *k*-mer loci reveal variation linked to secondary metabolite production. Together, our approach provides a functional framework to connect reference-free *k*-mer GWAS output to annotated, biologically interpretable pangenomic regions in a complex crop genome.

## 2 Results

We developed a pangenome-based *k*-mer interpretation framework and applied it to fruit metabolite variation in 343 *Capsicum annuum* accessions [22]. In this section, we first describe the pangenomic frame-work used to interpret trait-associated *k*-mers. Then we demonstrate the framework on three selected phenotypes.

### 2.1 A pangenome-based framework for localizing and interpreting significant *k*-mers

To overcome the limited interpretability of reference-free *k*-mer GWAS output, we developed a workflow that links trait-associated *k*-mers to genomic positions, annotations, and syntenic context within a graph-based *Capsicum* pangenome. The framework builds on existing PanTools functionality for pangenome construction, genome annotation, homology grouping, and extraction of genomic regions [20, 21], and we extended this functionality with a new function to localize significant *k*-mers directly in the pangenome graph.

Figure 1 summarizes the four inputs and eight analysis and interpretation steps of the workflow.

**Fig 1.**
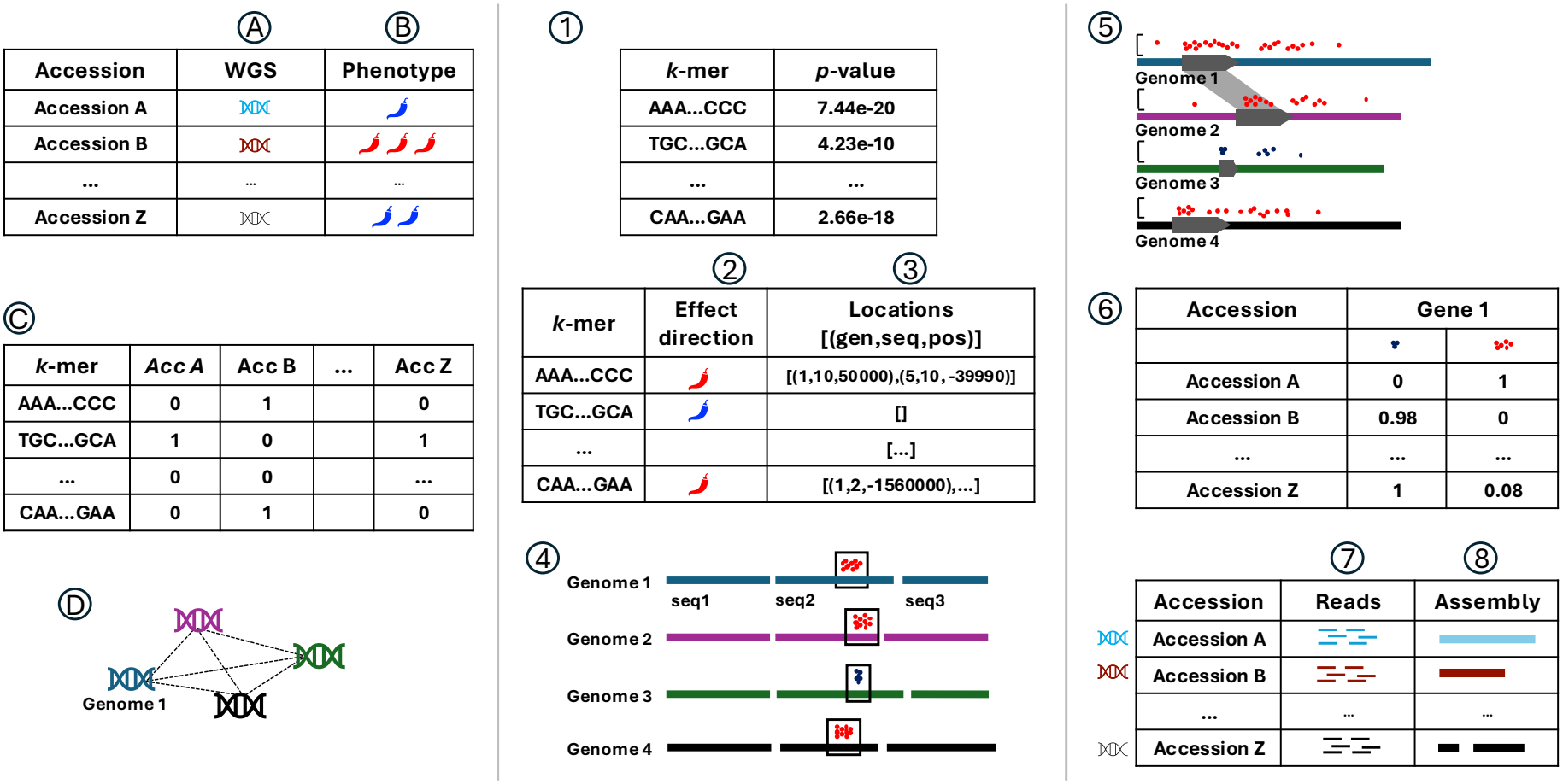
Workflow of pangenome-guided interpretation of *k*-mer GWAS signals. Required inputs are WGS data (A) and phenotypic data for sufficient accessions (B), a *k*-mer presence/absence table generated from the WGS data (C), a PanTools graph pangenome (D). To identify trait-associated loci, steps 1–4 are followed: *k*-mer GWAS to identify significant *k*-mers (1), classification of significant *k*-mers according to their association with higher or lower trait values (2), localization of these *k*-mers across all pangenome assemblies (3), and sliding-window aggregation of nearby locations into candidate loci (4). Several downstream analyses are possible: multi-genome Manhattan–synteny visualization of candidate loci (5), presence/absence-based haplotype analysis in resequenced accessions (6), ROI-specific read extraction and alignment to inspect locus variation (7), and assembly of the selected reads to reconstruct accession-specific haplotypes (8).

#### Inputs

A. Whole-genome resequencing (WGS) data for all accessions (*e*.*g*., accessions A–Z; see Section 5.1).
B. Phenotypic values for the same accessions (see Section 5.1).
C. *k*-mer table: a matrix of *k*-mer observations across accessions, represented as presence/absence or abundance. In this study, we used a filtered binary presence/absence (P/A) matrix generated from the resequencing data [7] (see Section 5.3).
D. Pangenome: a PanTools *k*-mer-based, de Bruijn graph pangenome constructed from genome assemblies and their annotations from the species or genus of interest (see Section 5.4).

#### Processing steps

1. *k*-mer GWAS: test *k*-mer variation across accessions for association with a phenotype and get significant *k*-mers (Inputs B and C; see Section 5.3).
2. *k*-mer effect-direction classification: use the P/A matrix and phenotypic values to determine whether each significant *k*-mer is associated with higher or lower phenotype values (Inputs B and C; see Section 5.7).
3. *k*-mer localization: use the pangenome graph (Input D) to retrieve the genomic coordinates of each significant *k*-mer in all genome assemblies in which it occurs. The *k*-mer size used for GWAS to be equal to or larger than the *k*-mer size used to construct the pangenome graph (Input D; see Section 5.5).
4. Locus identification: aggregate *k*-mer locations into candidate loci using a sliding-window approach (see Section 5.6).

#### Interpretation/visualization

5. Manhattan-synteny visualization: combine the association strength and effect direction of localized significant *k*-mers with gene annotations and syntenic relationships obtained through the pangenome to inspect each candidate locus across genome assemblies (see Section 5.8)
6. *k*-mer haplotype analysis: select sets of high- and low-associated *k*-mers belonging to a gene or subregion within the region of interest (ROI). Their presence/absence patterns across resequenced accessions are used to infer haplotype states and compare them with phenotype values (see Section 5.10).
7. ROI read selection: extract resequencing read pairs containing significant *k*-mers from the selected ROI and retain reads that align specifically to this region. The resulting alignments can be inspected in JBrowse 2 (see Section 5.9).
8. Accession-specific locus reconstruction: assemble the selected ROI reads to reconstruct the candidate locus in individual accessions and compare the resulting contigs with the corresponding (pan)genomic regions (see Section 5.9).

We applied the workflow to fruit metabolite variation in 343 *Capsicum annuum* accessions (Table S1) [22]. The accessions were sequenced (at approximately 25X coverage) and phenotyped for fruit metabolite levels at two locations. Location-adjusted BLUE values were then used as phenotypic input for association mapping. From the full metabolite dataset, we selected three representative traits for detailed pangenomic interpretation: total capsaicinoid content, dehydroxylinalool oxide, and *α*-ylangene (Table S1). These traits represent complementary dimensions of pepper fruit quality and metabolism. Total capsaicinoid content captures pungency, a defining sensory and breeding trait of pepper, whereas dehydroxylinalool oxide and *α*-ylangene represent aroma-related volatile terpenoids. Volatile profiles vary substantially among *Capsicum* genotypes, making these compounds useful contrasting traits for evaluating the framework beyond the well-characterized capsaicinoid pathway [23–25].

Reference-free *k*-mer analysis yielded 4,064,074,026 filtered *k*-mers, which were represented in a binary presence/absence matrix for *k*-mer GWAS (Section 5.3). For comparison with reference-based association mapping, variant calling against the *C. annuum* var. *annuum* ‘Zhangshugang’ reference genome [26] identified 1,439,631 high-quality SNPs used for SNP GWAS (Section 5.2). Finally, the *Capsicum* pangenome used for localization and interpretation of significant *k*-mers was constructed from 12 assemblies representing *C. annuum* and related *Capsicum* species (Table S2) and included gene annotations and homology information across the genomes.

### 2.2 Capsaicinoids: Resolving the haplotype variation at the *Pun1* locus

Pungency in *Capsicum* is mainly determined by capsaicinoid biosynthesis, with the *Pun1* locus being the major genetic determinant [27–29]. *Pun1* encodes capsaicin synthase, an enzyme essential for capsaicinoid production in the fruit placenta. Loss-of-function alleles, resulting from deletions and disruptive mutations, produce non-pungent/sweet peppers [27–29]. Previous genomic studies have further shown that the *Pun1* locus exhibits substantial structural variation, including copy-number variation of capsaicin synthase genes, across *Capsicum* genomes [5, 30]. Beyond *Pun1*, additional loci and genes affecting pungency include *Pun2*, which encodes a putative aminotransferase (*pAMT*), and *Pun3*, which encodes the R2R3-MYB transcription factor *CaMYB31* [30–32]. Total capsaicinoid content was calculated based on the BLUE values of capsaicin, dihydrocapsaicin, homocapsaicin, homodihydrocapsaicin, and nordihydrocap-saicin. We used this trait as a benchmark to test whether *k*-mer GWAS combined with pangenome-based localization could recover the established *Pun1* association and characterize associated sequence and structural variation beyond that resolved by reference-based SNP GWAS.

First, *k*-mer GWAS identified 9,611 significant *k*-mers for total capsaicinoid content, of which 5,430 were associated with higher capsaicinoid content and 4,181 with lower content based on their effectsize direction (Fig. 2A). Between 4,123 and 5,360 significant *k*-mers were localized in each *C. annuum* genome. Across all five *C. annuum* assemblies, chromosome 2 contained the highest density of associated *k*-mers (Fig. S1). Using the sliding-window approach, the localized significant *k*-mers were aggregated into an approximately 150 kb significant locus corresponding to the *Pun1* region and its tandem capsaicin synthase (CS) gene copies.

**Fig 2.**
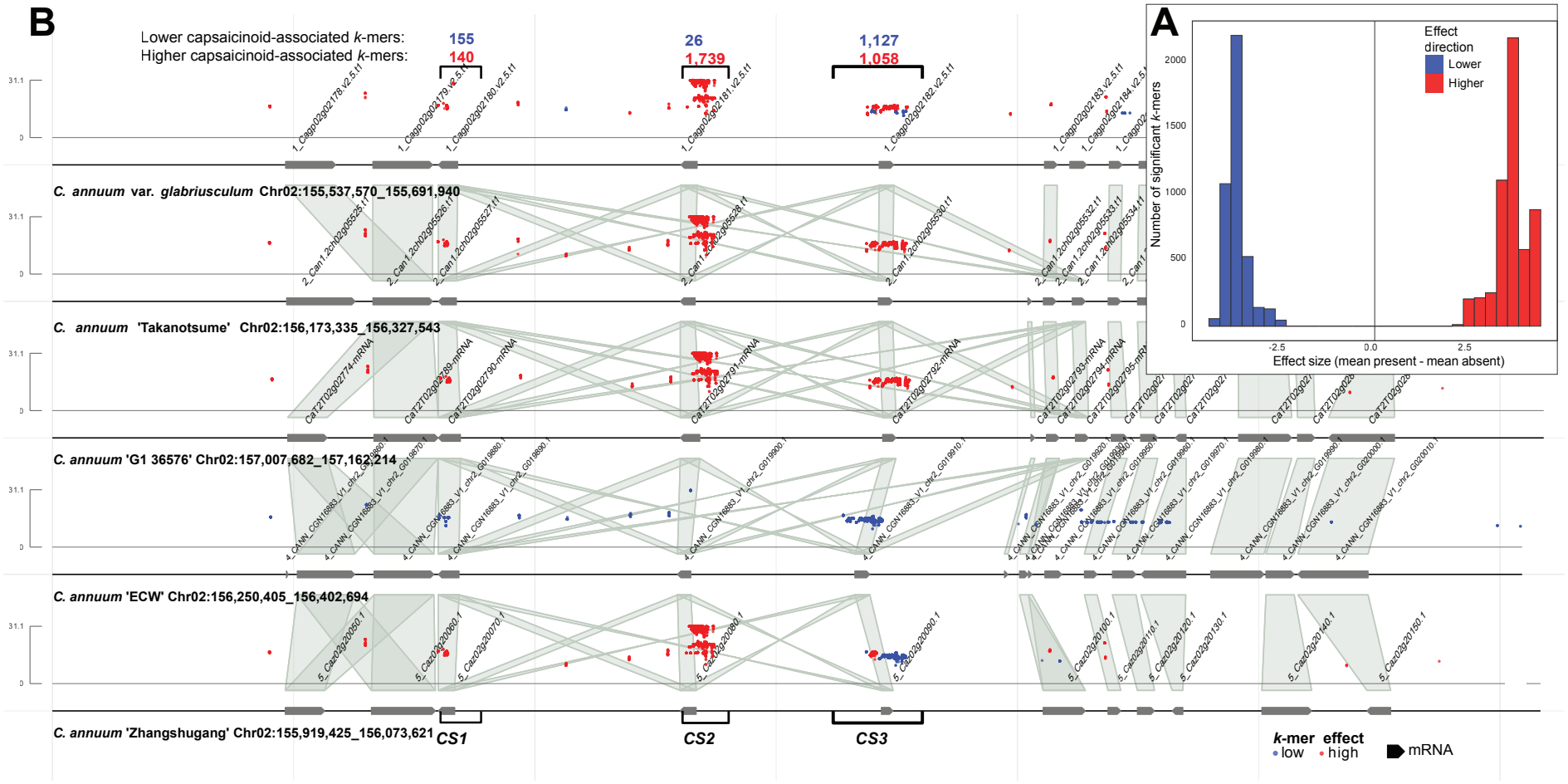
Capsaicinoids *k*-mer effect distribution and pangenome localization. Panel (A) Effect-size distribution of the 9,611 significant *k*-mers associated with total capsaicinoid content. Red indicates association with higher capsaicinoid content and blue indicates association with lower content. Panel (B) Combined manhattan–synteny plot showing the localization and association strength of significant *k*-mers across five *C. annuum* genome assemblies, together with gene annotations and homology relationships among the genomes. The three locations used to select *k*-mer sets associated with the *CS1, CS2*, and *CS3* genes are highlighted.

For comparison, SNP GWAS for total capsaicinoid content was performed against the pungent *C. annuum* var. *annuum* ‘Zhangshugang’ reference genome and identified a strong association signal in the region of the *Pun1* locus (Fig. S2). The main significant SNP region spanned Chr02:153,891,955-158,491,441 and contained 192 significant SNPs, with the most significant marker located at Chr02:155,951,872 (*p* = 7.0 *×*10^*−*26^). Both approaches therefore point to the same pungency locus. However, the pangenome-localized *k*-mers define an approximately 150 kb locus, compared with the approximately 4.6 Mb region identified by SNP GWAS (Fig. 2B; Fig. S2).

Notably, the localization and effect direction of significant *k*-mers differed strongly among the five *C. annuum* assemblies. Accessions ‘Takanotsume’ and ‘G1 36576’ contained only high-associated *k*-mers within the locus, whereas *C. annuum* var. *glabriusculum* and ‘Zhangshugang’ contained both high- and low-associated *k*-mers but were dominated by high-associated *k*-mers. In contrast, ‘ECW’ contained only low-associated *k*-mers, consistent with the low capsaicinoid content. These differences showed that no single assembly represented all trait-associated sequences within the locus, supporting the use of the pangenome for their localization (Fig. 2B).

Within this locus, significant *k*-mers were not evenly distributed but formed distinct clusters within and around the capsaicin synthase gene copies (Fig. 2B). Examination of the localized *k*-mers together with ROI-specific read alignments identified the known CS2/*Pun1* promoter and first-exon deletion and allowed its breakpoint to be resolved directly (Fig. S3). Approximately, 1,700 significant *k*-mers showed the same presence/absence pattern associated with this structural variant, including 26 low-associated *k* - mers spanning the deletion breakpoint (Table S3). This showed that groups of associated *k*-mers within the locus represent specific sequence and structural variants rather than independent association signals.

Following the locus-level inspection, we selected six non-overlapping *k*-mer sets representing high- and low-associated sequences around *CS1, CS2*, and *CS3* (Table S3). Their presence/absence patterns were examined across the 343 accessions and compared with the quantitative capsaicinoid phenotype (Table S4). The low-associated *CS2 k*-mer set showed the strongest correlation with capsaicinoid content (*r* =*−*0.82) and was also the most significant predictor in the linear regression analysis (*p* = 6.46*E−* 06) (Table S5). Also, occasional heterozygosity was detected. Finally, classification based on the *CS2* haplotype agreed with the sweet or hot phenotype for 91% of the accessions, supporting the major contribution of structural and sequence variation at *CS2* /*Pun1* (Table S4).

Combining the *CS1, CS2*, and *CS3* classifications produced seven of the 27 theoretically possible haplotype combinations (Table 1). No accession combined a hot-associated *CS1* haplotype with a sweet-associated *CS2* haplotype, consistent with the linkage pattern observed in the SNP data. The four common homozygous haplotype combinations occurred in both sweet and hot accessions, although their frequencies differed strongly between the phenotype groups. Rare heterozygous signals were also detected and supported by inspection of the read alignments (Fig. S3; Table S6). These results showed that the three CS haplotype classifications captured the major difference between sweet and hot accessions but are not sufficient to explain quantitative variation in capsaicinoid content. Accessions whose phenotype did not match the predominant haplotype pattern provide useful candidates for investigating additional sequence or regulatory variation within the locus.

**Table 1.** Pungency phenotypes across seven *k*-mer-based *CS1, CS2*, and *CS3* haplotype combinations. The numbers of sweet/non-pungent and hot/pungent accessions and their quantitative total phenotypic (Ph.) ranges are shown.

| **Haplotype** |  |  | **Phenotype** |  |  |  |
|---|---|---|---|---|---|---|
| ***CS1*** | ***CS2*** | ***CS3*** | **Sweet** | **Ph. range** | **Hot** | **Ph. range** |
| Sweet | Sweet | Sweet | 146 | 18.4–22.9 | 3 | 23.2–25.6 |
| Sweet | Hot | Sweet | 1 | 22.9 | 14 | 24.7–26.2 |
| Hot | Hot | Sweet | 3 | 22.1–22.9 | 1 | 26.0 |
| Hot | Hot | Hot | 22 | 19.1–22.9 | 149 | 23.0–26.4 |
| Hot | Hot | Heterozygous | 0 |  | 1 | 26.1 |
| Heterozygous | Hot | Heterozygous | 0 |  | 1 | 25.7 |
| Heterozygous | Heterozygous | Heterozygous | 1 | 19.8 | 1 | 25.3 |
| **Total** |  |  | **173** |  | **170** |  |

Together, these analyses showed that *k*-mer GWAS recovered the established *Pun1* association while pangenome localization narrowed the signal and connected it to assembly-specific sequence and structural variation. The CS2/*Pun1* deletion explained the strongest separation between sweet and hot accessions, while the combined *CS* haplotypes and phenotype exceptions indicated that additional variation within or around the locus may contribute to quantitative capsaicinoid content.

### 2.3 Dehydroxilinalool oxide

Specific terpene-derived aroma compounds provide a useful way to study how genetic variation affects downstream modifications in the terpenoid pathway. We investigated the genomic association of variation in dehydroxylinalool oxide (DHO), a linalool-derived monoterpenoid contributing to variation in fruit volatile profiles. Linalool and its oxygenated derivatives are products of terpenoid metabolism, a pathway shaped by expanded and functionally diverse terpene synthase (*TPS*) gene families in plants. Because *TPS* genes often occur in duplicated or structurally variable genomic contexts, dehydroxylinalool oxide provides a useful case to test whether pangenome-localized *k*-mers can move beyond a reference-genome association interval and help interpret candidate haplotypes in relation to gene content and homology [33, 34].

SNP GWAS for dehydroxylinalool oxide identified a single, clearly defined association peak above the genome-wide significance threshold on chromosome 10 of the *C. annuum* var. *annuum* ‘Zhangshugang’ reference genome (Fig. S4). The significant region spanned Chr10:38,292,723-40,679,226 and contained 55 significant SNPs, with the most significant marker located at Chr10:39,527,197 (*p* = 1.0 *×*10^*−*10^). Using *k*-mer GWAS, we identified 5,507 significant *k*-mers for dehydroxylinalool oxide, of which only 336 were associated with higher trait values, while 5,171 were associated with lower trait values based on effect-size direction (Fig. S5, S6). The significant *k*-mer set was therefore strongly enriched for *k* - mers associated with lower dehydroxylinalool oxide values. At the genome-wide level, the significant *k*-mers defined a single major locus on chromosome 10 that overlapped and was comparable in size to the interval identified by SNP GWAS; no additional major association locus was detected (Fig. S7). Pangenome localization revealed a strongly contrasting pattern across the five *C. annuum* assemblies. In the ‘Zhangshugang’ genome, the localized signal consisted almost exclusively of 340 high-associated *k*-mers, while ‘ECW’, ‘G1 36576’, ‘Takanotsume’, and *C. annuum* var. *glabriusculum* contained between 4,537 and 5,203 low-associated *k*-mers in the corresponding syntenic region (Fig. 3, S7).

**Fig 3.**
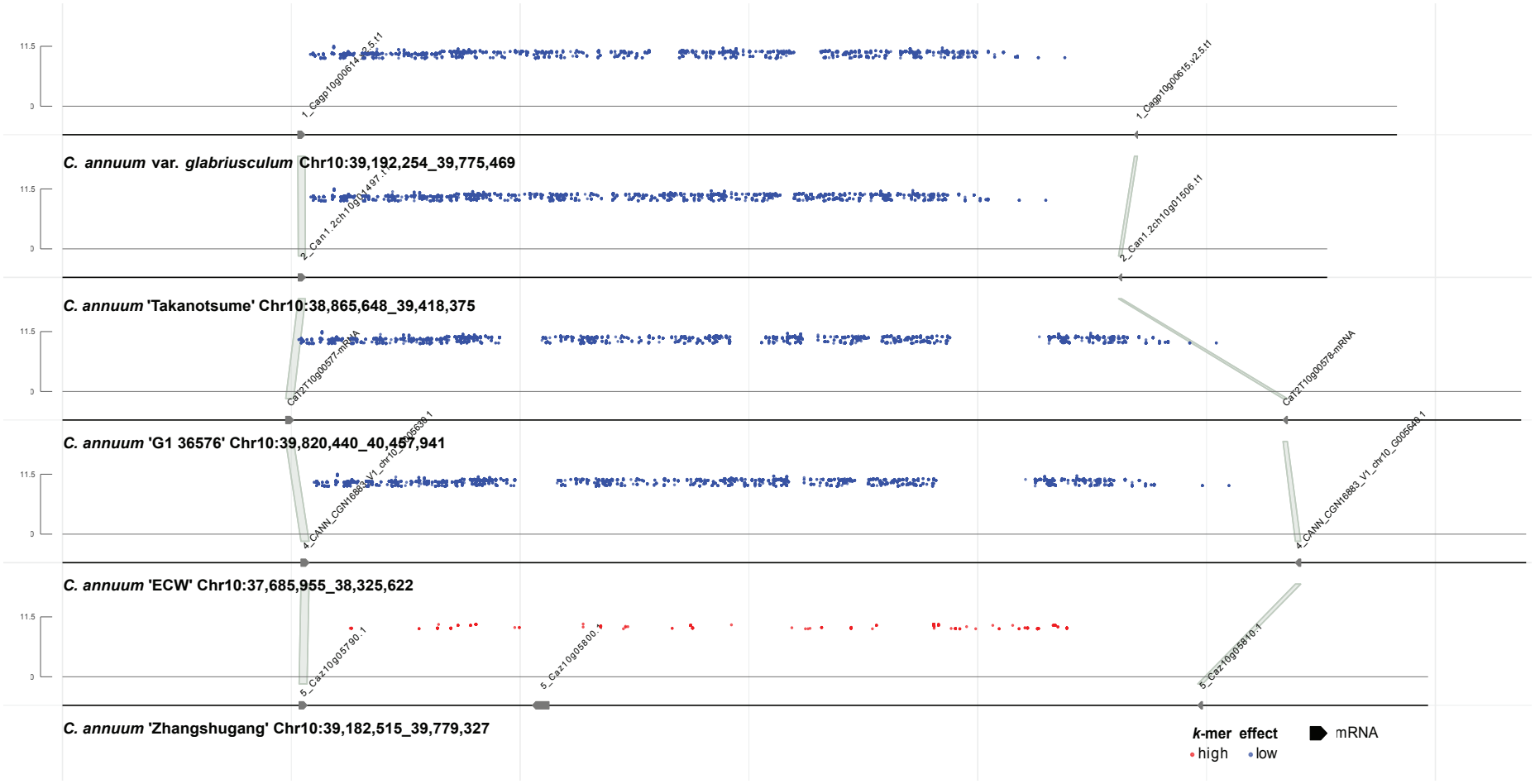
Dehydroxylinalool oxide *k*-mer effect distribution and localization. Manhattan and synteny plot for the five *C. annuum* genomes, including the significant *k*-mer localization and the gene homology in comparison to the other genomes.

Strikingly, the ‘Zhangshugang’ interval contained the terpene synthase gene *Caz10g05800*.*1*, suggesting that the association signal was linked to a distinct local *TPS* -associated haplotype (Fig. 3). However, pangenome-based homology grouping showed that this was not a simple presence/absence pattern of the *TPS* gene. Homologous *TPS* genes, with high similarity to *Caz10g05800*.*1*, were detected in all genomes, but were located approximately 10 Mb upstream of the associated interval rather than within the localized *k*-mer signal (Fig. S8, Table S7).

Because *k*-mers that are shared among these homologous gene copies are marked as present in all/many genomes the binary P/A analysis could not show whether an additional gene copy contributed to the high dehydroxylinalool oxide phenotype. The *k*-mer GWAS could instead distinguish the broader local haplotype containing *Caz10g05800*.*1*, while its contribution remained unresolved.

Based on the presence of *k*-mers in the 40 Kb up and downstream of *Caz10g05800*.*1* (Table S3), we know that only 29 (8.5%) of the accessions have the high-associated ‘Zhangshugang’ genotype, typically, but not exclusively associated with higher phenotypic scores (20.9 on average) (Table S8). To assess whether all accessions with this haplotype contain the gene, ROI-specific reads were extracted for six high and six low DHO accessions (Table S9). Read mapping showed that all six high DHO accessions had continuous coverage across *Caz10g05800*.*1*, implying that all accession with this haplotype contain the gene, whereas the six low DHO accessions showed no coverage across this gene (Fig. S9). When focusing on *Caz10g05800*.*1*, the absence of read alignments across the gene and its immediate up- and downstream regions in the low phenotype accessions further supported the absence of the gene and the variation of the locus compared to high phenotype accessions (Fig. S10). However, a few accessions that do not match the high-associated haplotype, do have a phenotypic score *≥*21 (Table S8), suggesting a different genetic mechanism to achieve high DHO production.

Together, the dehydroxylinalool-oxide case illustrated a different strength of the pangenomic workflow than the *Pun1* example. Rather than resolving a known deletion and sequence variation, pangenome-localized *k*-mers highlighted a *TPS* -associated haplotype and homology grouping showed that related *TPS* copies were present elsewhere in the pangenome, obliviating a *k*-mer signal on the gene itself. This allowed the association signal to be interpreted as a difference in local haplotype and gene-context variation, rather than as simple gene presence/absence. The result demonstrated how combining *k*-mer GWAS with pangenome localization, annotation, and homology information could refine candidate interpretation in duplicated metabolic gene families.

### *2*.*4 α*-Ylangene

Building on the analysis of monoterpene-derived aroma variation, we next investigated sesquiterpene compounds to explore how structural variation in terpene biosynthetic regions contributes to aroma diversity in *Capsicum*. This analysis focused on *α*-ylangene, a sesquiterpenoid volatile contributing to variation in fruit aroma profiles. Like other terpenoid metabolites, *α*-ylangene is expected to be influenced by terpene-synthase activity and by variation in local genomic regions containing duplicated or structurally variable metabolic genes. This trait therefore provided a third test case for the pangenome-localization frame-work, not focused on a single known causal gene, but on the interpretation of a broader and structurally complex association signal [34, 35].

First, SNP GWAS for *α*-ylangene identified a significant association on chromosome 9 in the *C. annuum* ‘Zhangshugang’ reference genome (Fig. S11). The significant region spanned chr09:12,589,880–15,196,341 and contained 281 significant SNPs, with the most significant marker located at chr09:12,590,933 (*p* = 1.3 *×*10^*−*11^). The size of this 2.61 Mb region suggested that the association was unlikely to be resolved by a SNP alone and motivated a pangenome-guided inspection of the surrounding haplotype structure.

Using *k*-mer GWAS, we identified 7,444 significant *k*-mers for *α*-ylangene, of which 5,387 were associated with higher trait values and 2,057 with lower trait values based on effect-size direction (Fig. S12). The significant *k*-mers identify a single 2.65 Mb trait associated locus on chromosome 9 (Chr09:12,580,590-15,229,919 in ‘Zhangshugang’), slightly broader than the locus identified by SNP GWAS. Across the five *C. annuum* assemblies, ‘Takanotsume’, *C. annuum* var. *glabriusculum*, and ‘Zhangshugang’ were dominated by high-associated *k*-mers, whereas ‘G1 36576’ is dominated by low-associated *k*-mers in the corresponding interval, and ‘ECW’ contains exclusively low-associated *k*-mers (Fig. 4, S13).

**Fig 4.**
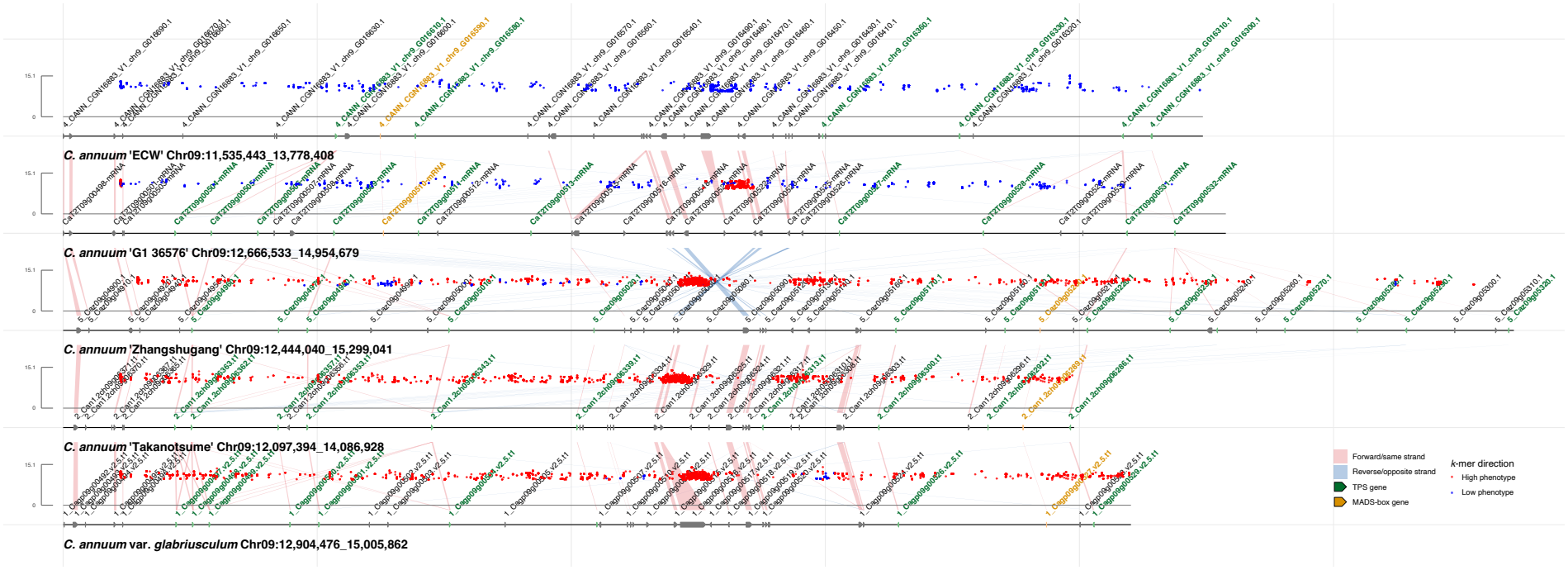
*α*-ylangene *k*-mer effect distribution and localization. Manhattan and synteny plot for the five *C. annuum* genomes, including the significant *k*-mer localization and the gene homology in comparison to the other genomes. Manhattan points show significant *k*-mers positioned within each genomic region and colored according to their phenotypic effect direction (red, higher phenotype; blue, lower phenotype). Red links connect genes on the same strand, whereas blue links connect genes on opposite strands. Terpene synthase (TPS) genes are highlighted in green and MADS-box genes in yellow.

Using synteny, we identified a large inversion of approximately 2 Mb. This rearrangement was not apparent from the SNP-GWAS peak alone, but became visible after localizing significant *k*-mers across multiple assemblies from the pangenome and comparing their gene order and orientation. In addition, pairwise alignment of the ‘Zhangshugang’ and ‘ECW’ genomes defined the exact transition boundaries of the inversion within the *α*-ylangene locus (Fig. S14). Interestingly, in ‘Zhangshugang’, the terpene synthase gene *Caz09g04960* was located in the collinear region approximately 138 kb upstream of the left inversion boundary. While no (TPS) gene was annotated at the corresponding position in ‘ECW’, TBLASTN identified strong exon matches at Chr09:11,759,641–11,762,232 and splice-aware alignment with miniprot supported a complete, intact protein-coding gene at this position, with 96.2% protein identity to *Caz09g04960*. This unannotated locus is therefore the most likely corresponding locus of *Caz09g04960*, despite more distant orthologs elsewhere. To compare *Caz09g04960* with previously reported *Capsicum* terpene synthases, we performed a BLASTP search against the proteins analyzed by Zhang et al. [33]. *Caz09g04960* showed 96.2% to a chromosome 9 protein designated TPS67 in that study. TPS67 was classified within the TPS-a subfamily, whose members commonly encode sesquiterpene synthases, but TPS67 itself was not functionally characterized.

Based on the pangenome, the distribution of high and low *k*-mers suggests that the organization of this locus (i.e. having the inversion or not) is strongly linked to the phenotype (producing *α*-ylangene or not). To investigate this relationship further, ROI-specific reads were extracted for six high- and six low *α*-ylangene accessions (Table S10) and visualized in JBrowse. We selected a 136 Kb structural variation just downstream of *Caz09g04960* with a consistent presence-absence pattern (Fig. S15), and a locus with a very high concentration of *k*-mers right in the middle of the inversion to characterize the haplotypes in the 343 resequenced accessions (Table S3). Table S11 shows that the majority of accessions (296) has the deletion and the 47 accessions with the insertion are concentrated in the high phenotypic range (with several exceptions), suggesting that the inversion might affect the expression of *Caz09g04960*. The same trend becomes apparent from the *k*-mers in the middle of the inversion, where accessions with a phenotypic score above 21 contain the high-associated *k*-mers. These two loci are insufficient for quantitative prediction, but they can be used to genotype the inversion and to predict the accessions witha high ( *≥*21) phenotype (92% correct label prediction). Moreover, they point out ‘interesting’ accessions that seem to escape the pattern, inviting more detailed locus analysis to understand the full regulation of this trait.

Taken together, the *α*-ylangene case illustrated the value of the pangenomic workflow for interpreting association signals in structurally complex regions. While the current analysis did not resolve a single causal gene or variant, pangenome-localized *k*-mers revealed that the association overlaps a large inversion-like haplotype on chromosome 9. This provided a more informative interpretation than the SNP GWAS interval alone and identified a defined structural region for future gene-level and functional follow-up.

## 3. Discussion

### 3.1 Pangenomic interpretation of *k*-mer GWAS

The combination of *k*-mer GWAS with a graph pangenome provided clear advantages for the localization and biological interpretation of association signals in a complex plant genome. We demonstrate the applicability of this approach in *Capsicum annuum*, a species with a large and repetitive genome and substantial structural variation. Significant *k*-mers could be localized within the pangenome and grouped into genomic regions across multiple assemblies, allowing association signals to be interpreted in relation to genome-specific coordinates, annotations, synteny and local haplotype structure.

In our examples, SNP and *k*-mer GWAS generally identified the same major genomic regions. This agrees with previous comparisons showing that *k*-mer associations commonly recover SNP-based associations, while also capturing structural variants and sequences that are missing from the reference genome [7]. The main value of our framework was therefore not simply the detection of additional loci, but the ability to connect associated *k*-mer clusters to SNPs, structural variants, and local haplotypes across multiple genome assemblies.

Recent studies have addressed the interpretation of *k*-mer associations using complementary analyses. Jaegle et al. localized resistance-associated *k*-mers using multiple wheat genomes, showing that broader genome representation improved locus detection and mapping compared with a single reference [13]. NodeGWAS instead performs association analysis directly with pangenome graph nodes, reducing marker redundancy while retaining graph context [19]. For polyploids, KMERIA uses *k*-mer abundance to account for allele dosage and subsequently maps significant *k*-mers to a graph pangenome [36]. Our framework is complementary to these approaches: it retains reference-free *k*-mer GWAS as the discovery step and uses the PanTools pangenome to connect significant *k*-mers to genome-specific coordinates, annotations, homology, synteny, and local haplotypes.

Beyond locus localization, clusters of significant *k*-mers with shared presence/absence patterns can be used to describe local haplotypes in resequenced accessions. At the *Pun1* locus, these clusters revealed existing haplotype combinations, including heterozygous configurations, while switches between high- and low-associated *k*-mer blocks can indicate recombination within an associated region. Even when a single causal variant cannot be resolved, this information helps prioritize haplotypes for downstream reconstruction, gene-level comparison, marker development, or even *de novo* sequencing and assembly.

Anticipating future application, this framework still holds several limitations that should be considered. The construction and analysis of large *k*-mer datasets remain computationally demanding, especially when many phenotypes are analyzed. In our analyses, pangenome construction, generation of the *k*-mer presence/absence table, and genome-wide association testing accounted for most of the runtime and required several days. Once these resources were available, localization of significant *k*-mers, locus aggregation, and ROI-based read selection generally required minutes to hours, depending on the number of *k*-mers, genomes, and reads analyzed. Approaches that reduce the number of *k*-mers that need to be tested, such as binning strategies (i.e. grouping *k*-mers with identical P/A patterns) or minimizer-based methods, may improve efficiency in future implementations, but careful considerations are needed to avoid the elimination of crucial variation. In addition, although the workflow helps localize association signals, the interpretation of association signals remains challenging for complex traits, where several loci or larger haplotypes may contribute to the phenotype.

The resolution of the analysis depends on the completeness and genotypic/phenotypic diversity of the underlying pangenome. Variation that is absent from the included assemblies cannot be represented, and structurally complex regions may still require long-read sequencing for local reconstruction and accurate interpretation. This was clear for *α*-ylangene, where the pangenome localized the signal to a candidate region that is inverted in some genomes, but did not yet resolve a causal gene or variant. However, in all use cases, interesting genotypes (e.g. escaping the pattern) were identified by the analysis pipeline. Finally, while the integration of *k*-mer GWAS and graph pangenomes offers substantial advantages for locus-level interpretation, further automation of the workflow and downstream (interactive) investigation/visualization would make the approach easier to reproduce and apply more broadly within the plant genomics community.

### 3.2 Application to *Capsicum annuum* and beyond

Understanding the genetic basis of metabolite diversity in *Capsicum* requires approaches that capture genomic variation beyond single-nucleotide polymorphisms. Here, we combined *k*-mer GWAS with a pangenomic interpretation to investigate the genomic architecture of selected secondary metabolites in*C. annuum*. Rather than only identifying associated markers, this approach allowed significant *k*-mers to be placed back into genomic, syntenic, and annotation context, while relevant haplotypes could be characterized.

In the three use cases under investigation *k*-mer GWAS typically identified the same locus as SNP GWAS, which is reassuring. The major advantage shows up in the locus analysis, because significant *k* - mers do not only capture sequence variation, but also structural variation. For instance, in the capsaicinoid case, the known CS2 promoter/first-exon deletion can be pinpointed precisely, not only in the assemblies, but also in the resequenced accessions. Also, unexpected heterozygosity was discovered through the *k*-mer approach.

The distinction between high- and low-associated *k*-mers is a powerful way to gain insight into factors that contribute positively or negatively to a trait. The presence of both high- and low-associated *k*-mers at a certain position might indicate heterozygosity (as seen in the capsaicinoid case), and the transitions between high- and low-associated *k*-mers, as visible in the *α*-ylangene case, may point to recombination within a trait-associated locus. This type of information is relevant for breeding applications, and can inform targeted experiments. It also emphasized the need to pangenomic localization, as a single genome typically contains only part of the *k*-mers, depending on its phenotype. Together with the locus visualization, contributing factors, which are hidden in identified haplotypes, can be identified. The terpene synthase in the dehydroxylinalool-oxide case is a good example of this. However, the *TPS* gene within the associated region remains a candidate that may contribute to dehydroxylinalool-oxide variation, but its effect cannot yet be separated from promoter, gene-context, or other linked variation within the haplotype.

In all three cases, groups of *k*-mers corresponding to genomic events can be used to quickly characterize the haplotype composition in the resequenced accessions (Table S4, S8, S11). This opens up the way toward genomic prediction, even though the small-scale setup in this study is insufficient for quantitative phenotype prediction. A more systematic approach across an entire associated locus might be able to explain the full regulation of a trait.

Together, these examples show that metabolite variation in *Capsicum* can involve known loci, homologous gene differences, and larger structural rearrangements, including structural variation that may not be captured by SNP-based approaches alone but can be detected using pangenome-localized k-mers. For breeding, this distinction is important as a regional SNP marker may be sufficient to track a strong locus such as *Pun1*, but traits influenced by duplicated genes or structural haplotypes may require markers that better represent the underlying haplotype. The *Pun1* analysis further showed that the low-associated *CS2 k*-mer set was more strongly correlated with capsaicinoid content than the corresponding high-associated set. This indicates that *k*-mers marking a low-phenotype haplotype can sometimes provide stronger predictors than those associated with the high phenotype. Pangenome-localized *k*-mers can therefore help prioritize candidate regions and identify informative markers for metabolite traits, especially when the relevant variation is not well represented by a single reference genome.

Although demonstrated in *Capsicum*, this framework is broadly applicable to other crop species and population genomics studies, particularly where structural variation and gene presence-absence variation play a major role in trait diversity. Recent work in crop pangenomics and association mapping high-lights the importance of incorporating genetically diverse genome assemblies to capture trait-associated variation beyond a single reference [3–5, 37].

Our approach may be especially useful in breeding contexts where the target trait is influenced by complex haplotypes that are not adequately represented by individual markers from SNP-based GWAS. For traits such as pungency or fruit aroma, pangenome-localized *k*-mers could support the development of markers that better represent causal or linked sequence variation across diverse germplasm. This may improve marker transferability across genetic backgrounds, particularly when important alleles are missing from, rearranged relative to, or poorly represented in a single reference genome.

The framework may also be relevant for polyploid species, where conventional SNP-based GWAS is complicated by genome structure, multi-allelic variation, and allele dosage effects. Recent work with KMERIA showed that *k*-mer copy-number information can be used for dosage-aware association mapping in complex polyploids and combined with a graph pangenome for subsequent localization [36]. PanTools can construct and query pangenomes containing both diploid and polyploid genomes, and the localization step itself is not restricted by ploidy. The workflow can therefore be combined with the *k*-mer association strategy most appropriate for the population, including presence/absence-based approaches or abundance-based methods that account for allele dosage. Future developments will likely focus on improving computational efficiency, expanding graph pangenome resources, and integrating *k*-mer association signals with functional annotation, expression data, and breeding-oriented marker design.

## 4. Conclusions and future work

Our study demonstrates that interpreting the results of *k*-mer GWAS using a *k*-mer graph pangenome provides an effective framework for identifying and interpreting the genomic basis of metabolite variation in *Capsicum annuum*. By enabling the detection and localization of structural and haplotype-level variation beyond conventional SNP analyses, the approach improves candidate gene discovery and facilitates the development of more informative breeding markers. As graph pangenomes continue to expand and computational methods become more efficient, this framework has strong potential to become a broadly applicable strategy for dissecting complex traits across diverse crop species. Future research should focus on the biological validation of candidate loci and haplotypes identified in this study using gene expression analyses, long-read sequencing, and gene-editing approaches. In parallel, graph pangenome resources can be expanded by incorporating more genetically diverse *Capsicum* accessions and wild relatives, together with phased, chromosome-scale genome assemblies that more accurately represent structurally complex regions. Methodological developments should aim to improve computational efficiency through strategies such as *k*-mer binning, minimizer-based approaches, and hierarchical association methods to better exploit increasingly large genomic datasets. Standardized visualization and analytical tools will also be needed to facilitate the integration of association signals with graph genomes and haplotype reconstruction. Finally, combining *k*-mer association signals with transcriptomic, metabolomic, epigenomic, and functional annotation data will improve the prioritization of causal variants and accelerate the translation of informative *k*-mers and haplotypes into robust diagnostic markers for breeding applications.

## 5. Materials and methods

### 5.1 *C. annuum* data sources and phenotyping

Whole-genome sequencing (WGS) was conducted for 343 *Capsicum annuum* accessions. Genomic DNA was extracted and used to construct short-insert libraries according to the DNBSEQ standard protocol. Paired-end sequencing (2 × 150 bp) was performed using the DNBSEQ platform. Raw reads were processed with SOAPnuke v2.1.9 (BGI) [38] to remove adapter contamination, low-quality reads, and sequencing artifacts. Adapter sequences were trimmed if at least 25% of a read matched the adapter sequence, allowing up to two mismatches. Reads that were shorter than 150 bp after trimming were discarded and reads containing at least 1% ambiguous bases (N), polyX stretches longer than 50 bp, or at least 30% bases with Phred quality scores below 20 were removed. Only reads that passed all filtering criteria were retained for downstream analyzes.

Large-scale metabolite data were measured from fruits that were grown at two locations. Best linear unbiased estimates (BLUEs) (Table S1) for the study phenotypes were obtained by fitting a linear model with accession and trial included as fixed effects using `lm` function from the R package `stats` v4.2.1 https://www.R-project.org/. Estimated marginal means were calculated using the `emmeans` function from the emmeans package v1.6.2-1 [39].

### 5.2 SNP GWAS

Reads that passed all the filtering steps were aligned to the *C. annuum* var. *annuum* ‘Zhangshugang’ reference genome [3] using the NVIDIA Clara Parabricks implementation of BWA-MEM (v4.3.0) [40]. Read-group information was assigned per accession, and duplicate reads were marked during alignment. Variant calling was conducted per sample using Parabricks HaplotypeCaller in GVCF mode [41]. Individual gVCFs were combined, and joint genotyping was performed using the GATK 4.5 germline short variant discovery pipeline to obtain a multi-sample VCF containing SNPs and indels. Variant normalization and decomposition were performed using `bcftools norm` [42], removing duplicate records and left-aligning variants relative to the reference genome.

Before conducting the genome-wide association study (GWAS), variants with genotype quality (GQ) below 20 were set to missing, and markers with more than 5% missing data were discarded. All subsequent filtering was performed using PLINK v2.00a3 SSE4.2. Genotype data were filtered in three steps. First, markers were filtered using thresholds of missingness *≤*0.1, heterozygosity *≤*0.1, and minor allele frequency (MAF) *≥*0.05, and only biallelic SNPs were retained. Second, samples with missingness or heterozygosity greater than 0.1 were removed. Marker-level filtering was then repeated using the same thresholds. Missing genotypes were imputed using Beagle v5.4 with default parameters [43]. To reduce redundancy caused by strong linkage disequilibrium, variants were pruned in 1 Mb windows using an *r*^2^ threshold of 0.98, retaining one representative marker from each high-linkage disequilibrium cluster.

A kinship matrix and the first three principal components (PCs) of the genome-wide principal component analysis were estimated from the filtered and linkage-disequilibrium-pruned SNP dataset using PLINK v2.00a3 SSE4.2. Association analysis was performed using a linear mixed model in GEMMA v0.98.5 [44], with the first three PCs included as fixed-effect covariates and kinship included as a random effect. Marker associations were tested using the Wald test. GEMMA was run using the options `-miss 1 -maf 0 -hwe 0 -lmm 1`; input, kinship, covariate, and output files were specified separately for each phenotype. Genome-wide significance was determined using a Bonferroni-corrected threshold calculated as *p* = 0.05*/N*, where *N* was the number of markers tested, corresponding to *−*log_10_(0.05*/N*) in the Manhattan plots.

### 5.3 k -mer Genome-Wide Association Study (GWAS)

*k*-mer–based association analyses were performed as described in Voichek and Weigel [7]. A *k*-mer length of 31 bp was used following the default settings of the reference-free GWAS pipeline. For each accession, *k*-mers were counted directly from the filtered paired-end reads using KMC v3.2.4 [45]. Canonical *k*-mers were generated by collapsing each sequence with its reverse complement into the lexicographically smaller form and were retained if observed at least twice per sample to minimize *k*-mers derived from sequencing errors. Simultaneously, non-canonical *k*-mers were counted to recover strand information, which was assigned using `kmers_add_strand_information`. To limit the inclusion of rare variants, only *k*-mers present in at least five accessions and in at least 20% of samples were retained. A binary presence/absence matrix was generated using `build_kmers_table`. The kinship matrix was calculated from the *k*-mer matrix using `emma_kinship_kmers`, considering only *k*-mers with a minor allele frequency of at least 0.05.

Association testing for each phenotype was performed separately using GEMMA (v0.98.5) [44]. For each *k*-mer, a linear mixed model incorporating the *k*-mer–derived kinship matrix was fitted. Significance thresholds were determined using 100 phenotype permutations per trait. Next, the resulting significant *k*-mers were ranked by association score, and those in the top 10th percentile were selected as significant and were retained for downstream analysis. Samples lacking phenotype data were excluded from the respective analyses.

### 5.4 Graph pangenome construction

Genome assemblies and annotations of 12 *Capsicum* accessions (five *Capsicum annuum* and seven repre-sentatives of other *Capsicum* species) were collected to construct a genus-level pangenome. Contigs shorter than 5 kb were removed, and only the longest protein-coding isoforms (*>*30 amino acids) were retained using the PanTools QC pipeline (https://github.com/PanUtils/pantools-qc-pipeline). Genomes were oriented based on their longest alignment to the *C. annuum* ‘Zhangshugang’ reference. The pangenome was constructed using PanTools v4.3.3 [20, 21], incorporating structural and functional annotations. Homology groups were inferred using the `optimal_grouping` function with the `solanales_odb10` lineage, and proteins were clustered using a relaxation parameter of 5. For the *α*-ylangene ROI analysis, the relaxation parameter was set to 3.

### 5.5 Mapping significant *k*-mers to genomic coordinates

Significant *k*-mers identified by *k*-mer GWAS were looked up in the PanTools pangenome graph as exact sequence matches, considering both the forward sequence and its reverse complement. As each sequence in the graph is linked to its occurrences in the constituent genomes, a single graph lookup retrieves the genomic coordinates of each *k*-mer in all genomes in which it is present.

The lookup procedure can be applied to *k*-mers of any length equal to or larger than the *k*-mer size used to construct the PanTools database. When a query *k*-mer was longer than the graph *k*-mer size, its complete sequence was matched by following the corresponding path through one or more graph nodes. Only exact matches of the complete query sequence were retained. PanTools version 4.3.3, abbrev. commit ID dc4b3ff was used to run the function `kmer_localization`. The resulting graph-derived coordinates were validated against alignments of the significant *k*-mers to the corresponding linear genome assemblies. The significant *k*-mer alignments to the linear genome assemblies were done using bowtie v1.3.1 [46].

### 5.6 Aggregating localized *k*-mers into candidate loci

Localized significant *k*-mers were aggregated into candidate loci separately for each genome using a sliding-window approach. A window was retained when the number of significant *k*-mers within it was equal to or greater than a user-defined threshold. Overlapping or directly adjacent significant windows were then merged into larger loci. The window size, step size, and minimum *k*-mer threshold were selected according to the density and scale of the association signal for each trait. For capsaicinoids, we used a 2,000-bp window, a 500-bp step, and a threshold of 15 *k*-mers. For *α*-ylangene, these parameters were 50,000 bp, 10,000 bp, and 10 *k*-mers. For dehydroxylinalool oxide, they were 15,000 bp, 5,000 bp, and five *k*-mers. The resulting candidate loci were used for multi-genome visualization and subsequent ROI selection.

### 5.7 Effect-direction classification and *k*-mer prioritization

The direction of each significant *k*-mer association was determined by calculating the difference in mean phenotype between accessions in which the *k*-mer was present and those in which it was absent (mean present *−* mean absent). Positive values indicated an association with higher trait values (high-associated), while negative values indicated an association with lower trait values (low-associated). All significant *k*-mers had a defined, non-zero mean phenotype difference.

For downstream analysis and visualization, significant *k*-mers were separated according to their effect direction and ranked within each group by decreasing statistical significance ( *−* log_10_(*p*)). The absolute mean phenotype difference was used as a secondary ranking criterion. The top X high-associated and top X low-associated *k*-mers were subsequently selected.

### 5.8 Single and multi-genome visualization

Genome-wide SNP associations were visualized as Manhattan plots along the ‘Zhangshugang’ reference genome. For *k*-mer GWAS, significant *k*-mers were plotted at their localized positions in each of the five *C. annuum* genome assemblies. Association strength was represented by *−*log_10_(*p*), while color indicated association with higher or lower trait values.

For candidate loci, the *k*-mer Manhattan plots were combined with multi-genome synteny visualizations. This allowed the position, association strength, and effect direction of localized *k*-mers to be compared with gene annotations, homologous genes, and syntenic relationships across the genome assemblies.

Presence/absence heatmaps were used to summarize the distribution of selected high- and low-associated *k*-mers across accessions. Accessions were ordered by phenotype, while *k*-mers were ordered by effect direction and association rank. All visualizations were generated using custom Python and R scripts.

### 5.9 ROI definition and accession-specific locus reconstruction

Selected candidate loci were defined as regions of interest (ROIs) for detailed analysis and accession-specific reconstruction. The corresponding sequences were extracted from the pangenome assemblies using the `extract_region` function of PanTools v4.3.4 (branch `add_gfa_export`, commit 15d7e3d). Resequencing read pairs containing significant *k*-mers localized within each ROI were extracted and aligned to the corresponding genome assemblies using BWA-MEM [40], and only reads mapping exclusively to the ROI were retained (Chapter 6 in [47]). The resulting alignments were visualized to inspect read support and haplotype differences among accessions. For accession-specific reconstruction, the retained reads were subsequently assembled using MEGAHIT v1.2.9 [48], and the resulting contigs were compared with the corresponding pangenomic regions.

### 5.10 Haplotype analysis based on *k*-mers

Within the capsaicinoid ROI, six *k*-mer sets were defined to characterize variation around the three main capsaicin synthase genes, *CS1, CS2*, and *CS3*, and their immediate flanking regions (Table S3). For each gene, the significant *k*-mers associated with high and low capsaicinoid values were selected from their positions on the five *C. annuum* genome assemblies. The *k*-mers occurring in more than one set were excluded, resulting in six non-overlapping, gene- and haplotype effect-specific *k*-mer sets. For every resequenced genotype, we calculated the fraction of *k*-mers present for each set (Table S6). Based on these scores the haplotype was labeled as ‘Sweet’ ( *≥* 0.5 of low *k*-mers and *<* 0.5 high *k*-mers), ‘Hot’ (*≥ 0*.*5* of high *k*-mers and *<* 0.5 low *k*-mers), or else ‘Heterozygous’. The same methodology was applied in the other use cases (Table S3, S8, S11).

## Acknowledgments

We thank the G2P-SOL consortium for developing and characterizing the broader *Capsicum* population used in this study. This work was funded by the Top consortium for Knowledge and Innovation (TKI) programs LWV2004 ‘Pangenome architecture and differentiation of agronomical traits in the Capsicum clade’ and LWV23099 ‘From Omics data to novel breeding targets in pepper’.

## Data and code availability

Whole-genome resequencing data for the 343 *C. annuum* accessions will be deposited in a public sequence repository upon peer-reviewed publication. Accession numbers for all publicly available genome assemblies used in this study are provided in Table S1. PanTools, including the *k*-mer localization functionality, is available via bioconda (version 4.3.3, commit ID dc4b3ff). Custom processing and visualization scripts will be made available upon peer-reviewed publication.

## Notes

### Competing Interest Statement

The authors have declared no competing interest.

