## Supplementary figures for "Pangenome-guided interpretation of *k*-mer GWAS in *Capsicum*"

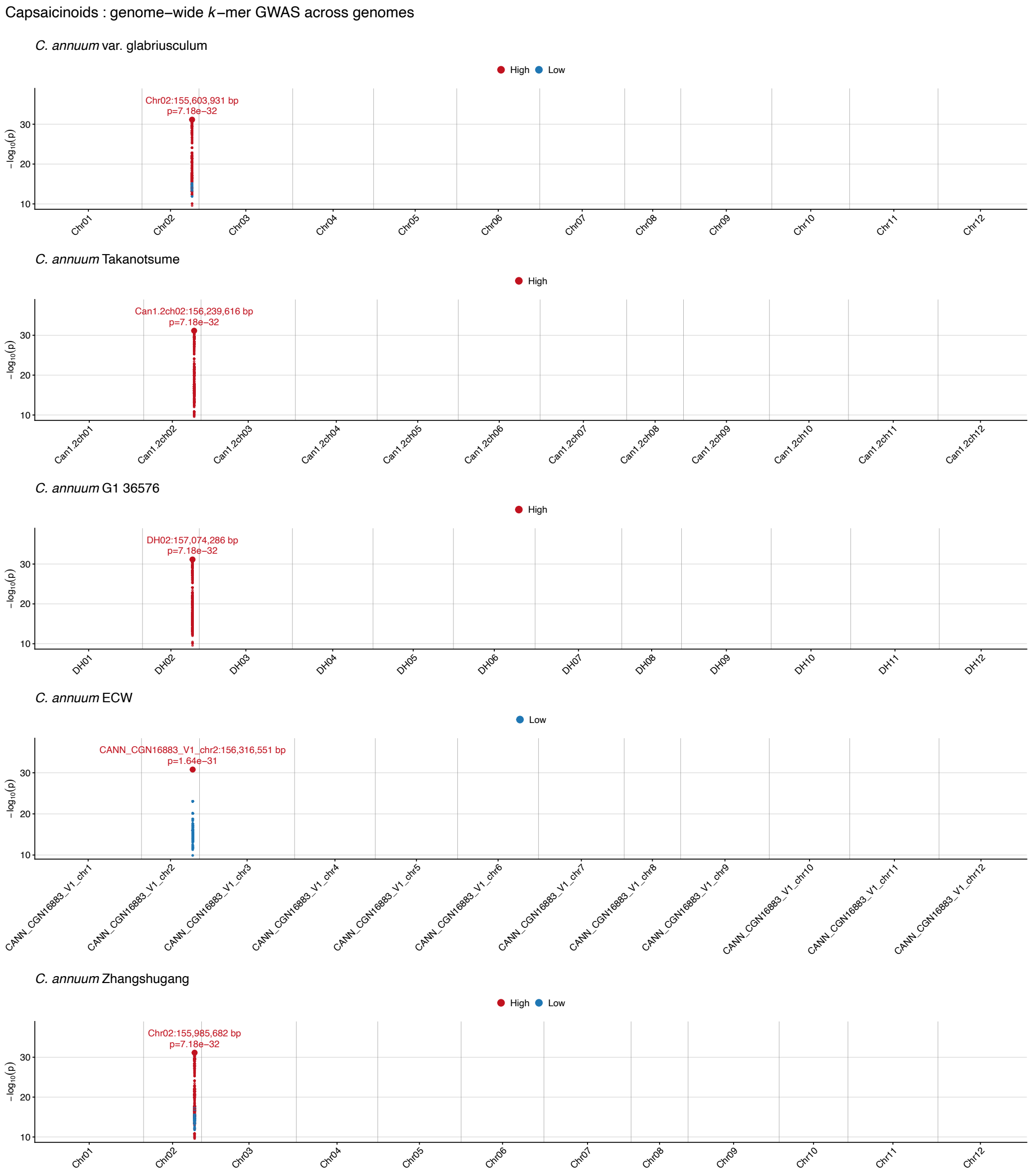


Figure S1: **Genome-wide *k*-mer association results for capsaicinoids.** Manhattan plots of *k*-mer genome-wide associations (GWAS) across the *C. annuum* genomes. Only significant *k*-mers are shown. The red highlighted *k*-mer positioned at the distal end of chromosome 2 marks the most significant association.

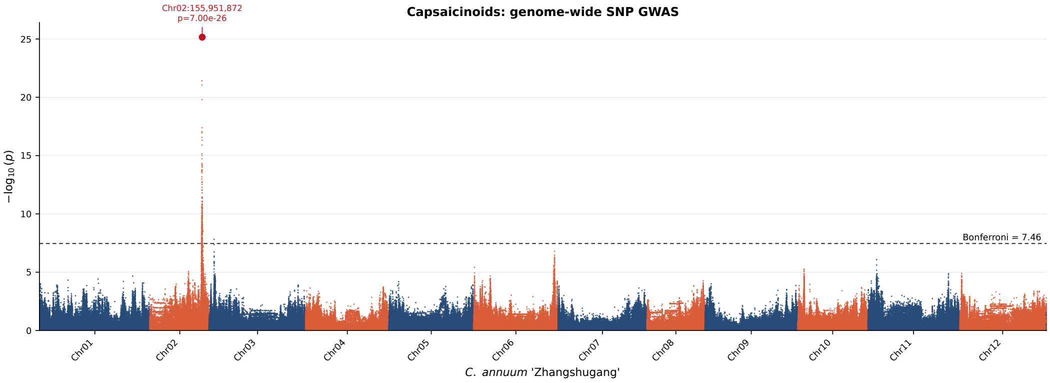


Figure S2: **Genome-wide SNP association results for capsaicinoids.** Manhattan plot of SNP genome-wide associations (GWAS) across the *C. annuum* ‘Zhangshugang’ genome. The dashed line indicates the Bonferroni-corrected genome-wide significance threshold ($-\log_{10}(p) = 7.46$), calculated from the number of SNP markers. The SNP highlighted in red on chromosome 2 represents the marker with the strongest statistical association.

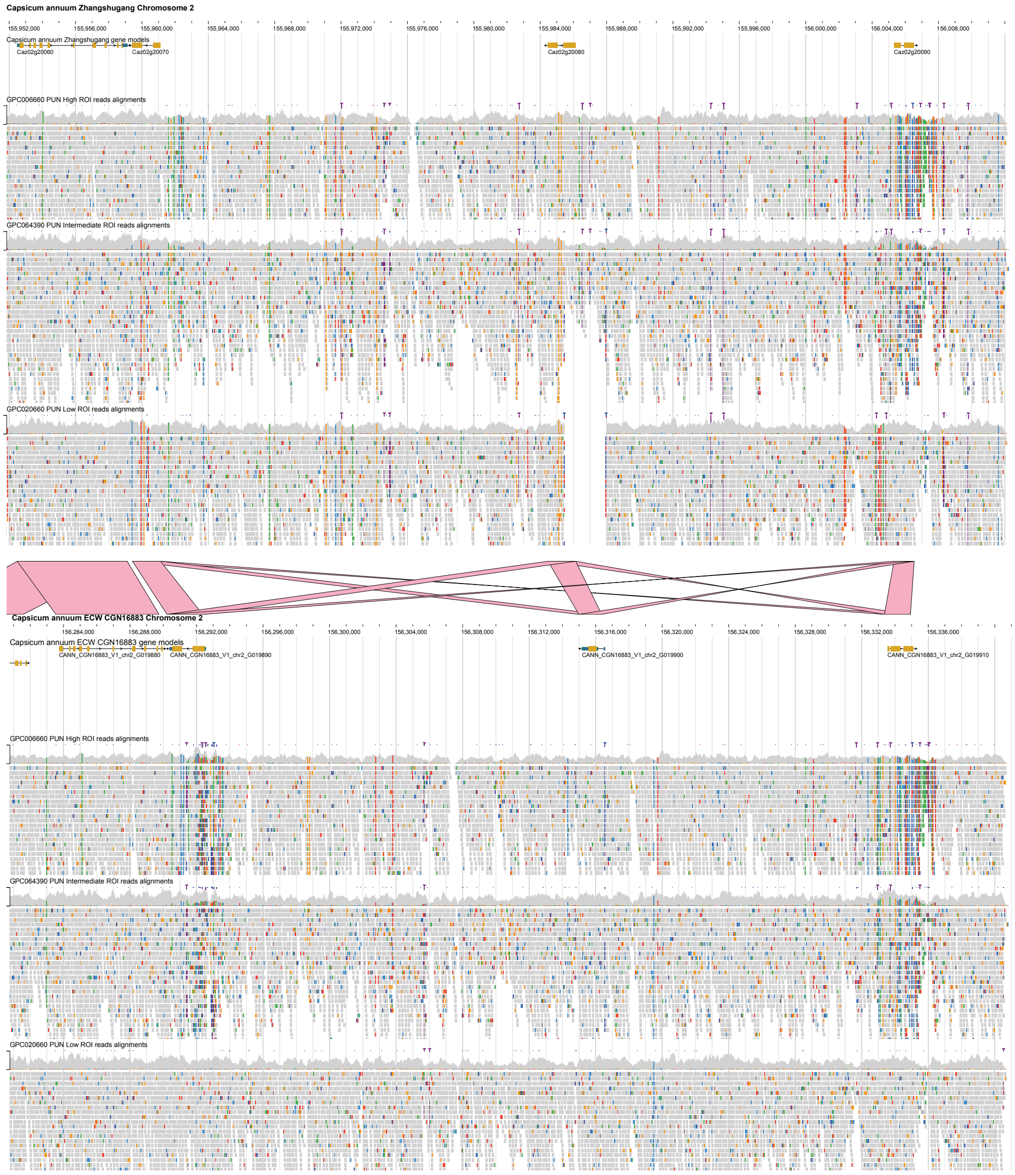


Figure S3: **Read alignments and synteny across the *CS1*, *CS2*, and *CS3* genes at the *Pun1* locus.** JBrowse 2 linear synteny view of the *Pun1* regions in the *C. annuum* ‘Zhangshugang’ (top) and ‘ECW’ (bottom) genomes. Pink bands connect genes assigned to the same homology groups. ROI-selected short reads are shown for accessions GPC006660 (pungent), GPC064390 (intermediate phenotype), and GPC020660 (low pungency). Grey blocks and shading represent aligned reads and their coverage, while colored marks indicate sequence differences relative to the corresponding genome.

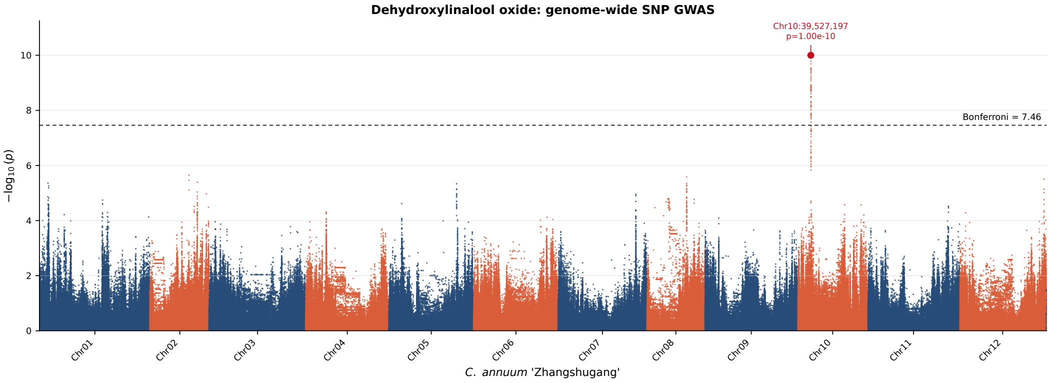


Figure S4: **Genome-wide SNP association results for dehydroxilinalool oxide.** Manhattan plot of SNP genome-wide associations (GWAS) across the *C. annuum* 'Zhangshugang' genome. The dashed line indicates the Bonferroni-corrected genome-wide significance threshold ($-\log_{10}(p) = 7.46$), calculated from the number of SNP markers. The SNP highlighted in red on chromosome 2 represents the marker with the strongest statistical association.

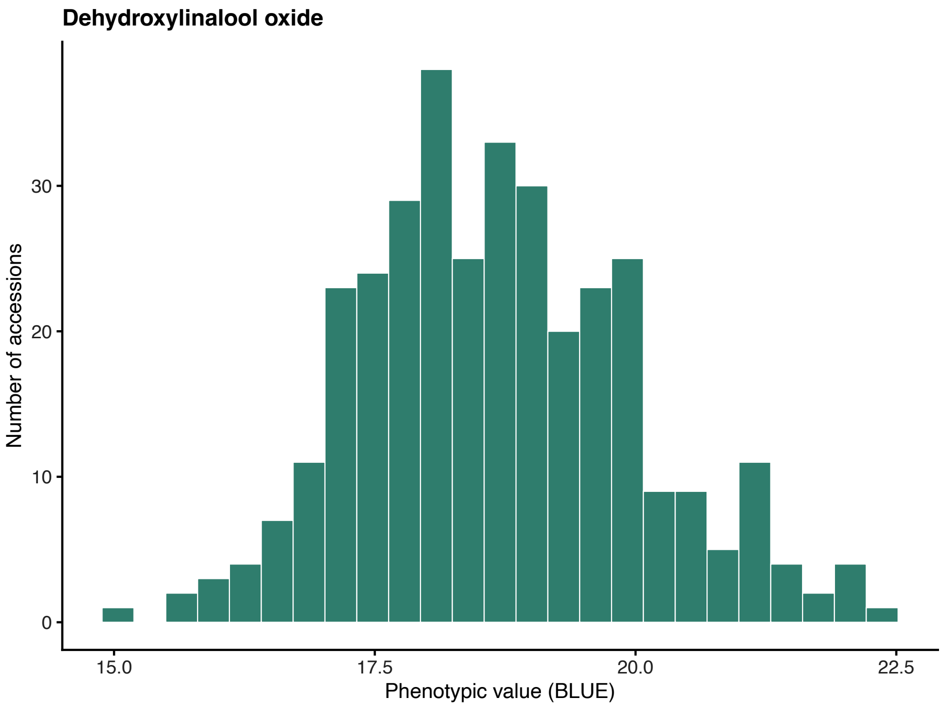


Figure S5: **Phenotypic distribution of dehydroxylinalool oxide.** Histogram showing the distribution of dehydroxylinalool oxide phenotype values across the 343 *C. annuum* accessions.

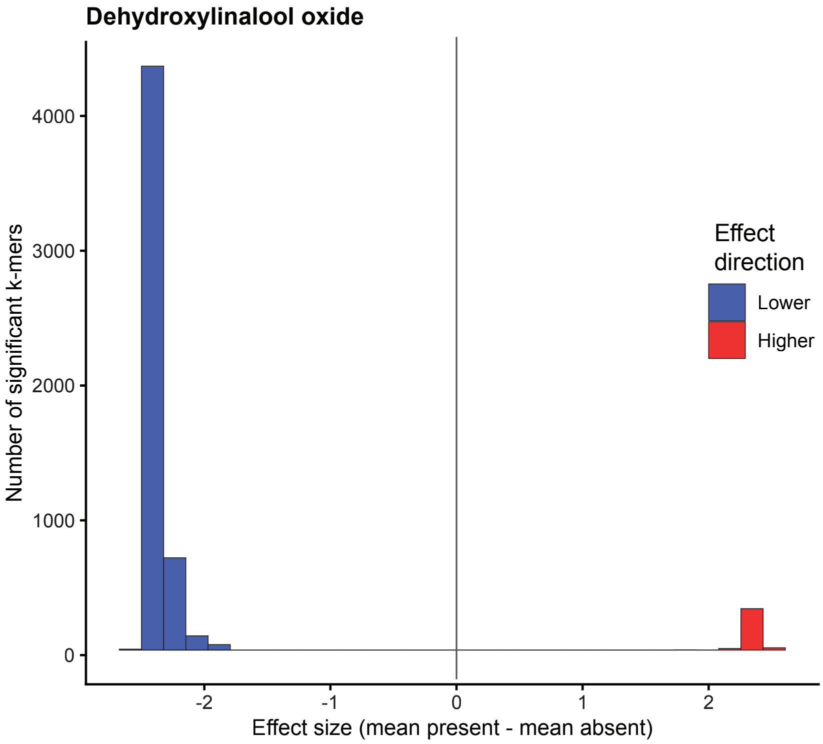


Figure S6: **Effect size distribution of significant *k*-mers associated with dehydroxilinalool oxide.** Histogram showing the difference in mean phenotype between accessions in which each significant $k$-mer was present and absent. Positive values indicate association with higher phenotype values, whereas negative values indicate association with lower phenotype values.

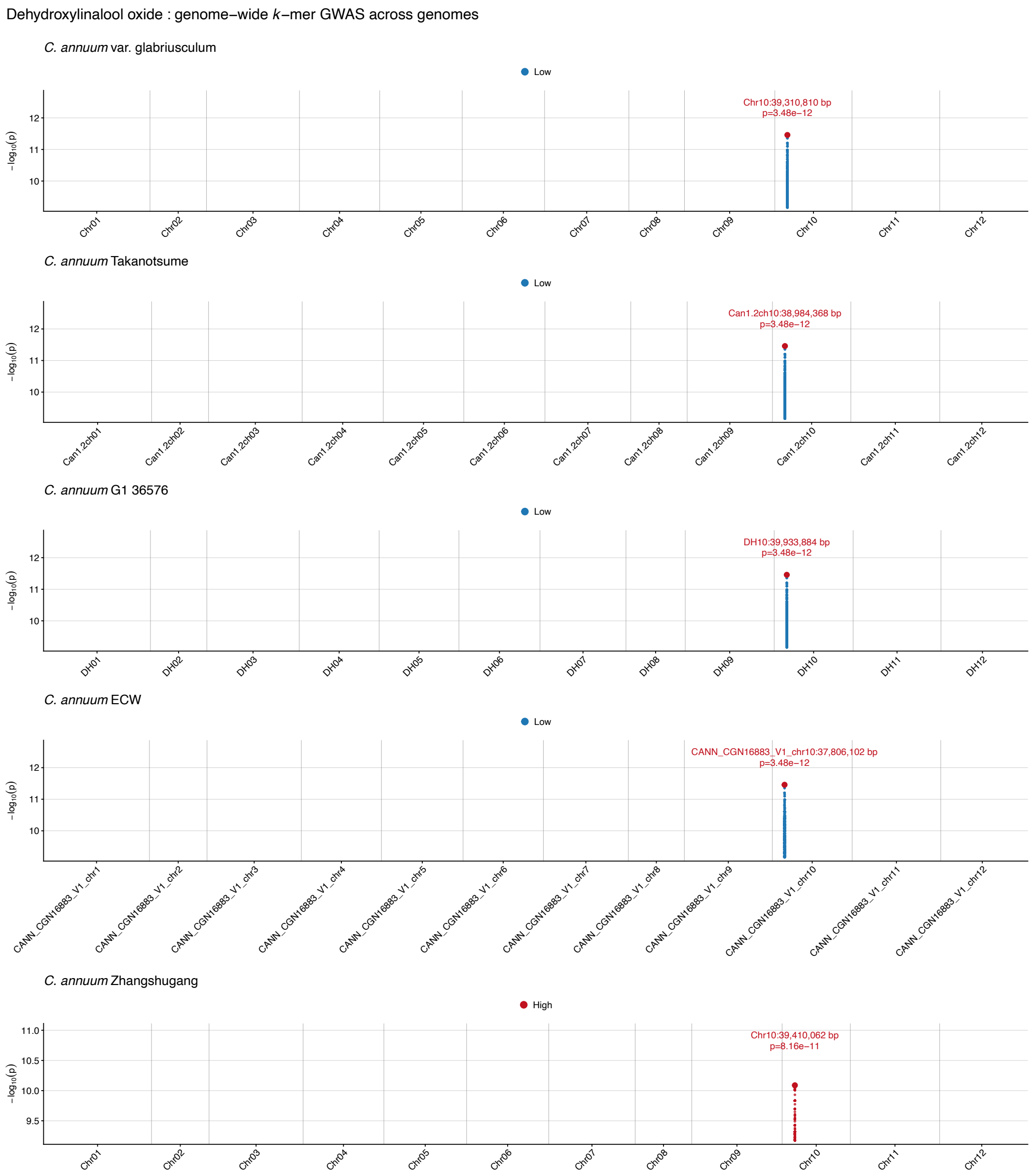


Figure S7: **Genome-wide *k*-mer association results for dehydroxilinalool oxide.** Manhattan plots of *k*-mer genome-wide associations (GWAS) across the *C. annuum* genomes. Only significant *k*-mers are shown. The red highlighted *k*-mer marks the most significant association.

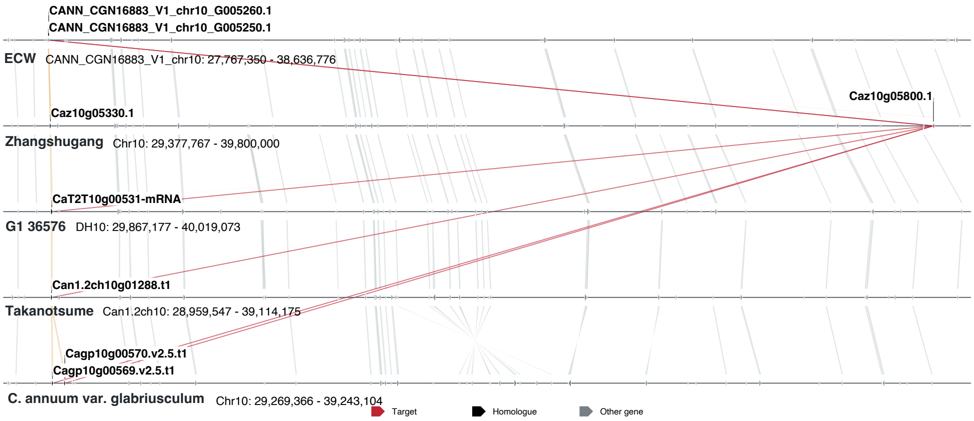


Figure S8: **Overview of the *Cza10g05800.1* homologs in the five *C. annuum* genomes.** Synteny plot of the five *C. annuum* genomes including the gene homology in comparison to the other genomes. The homology connections of the *Cza10g05800.1* gene homologs are highlighted in red.

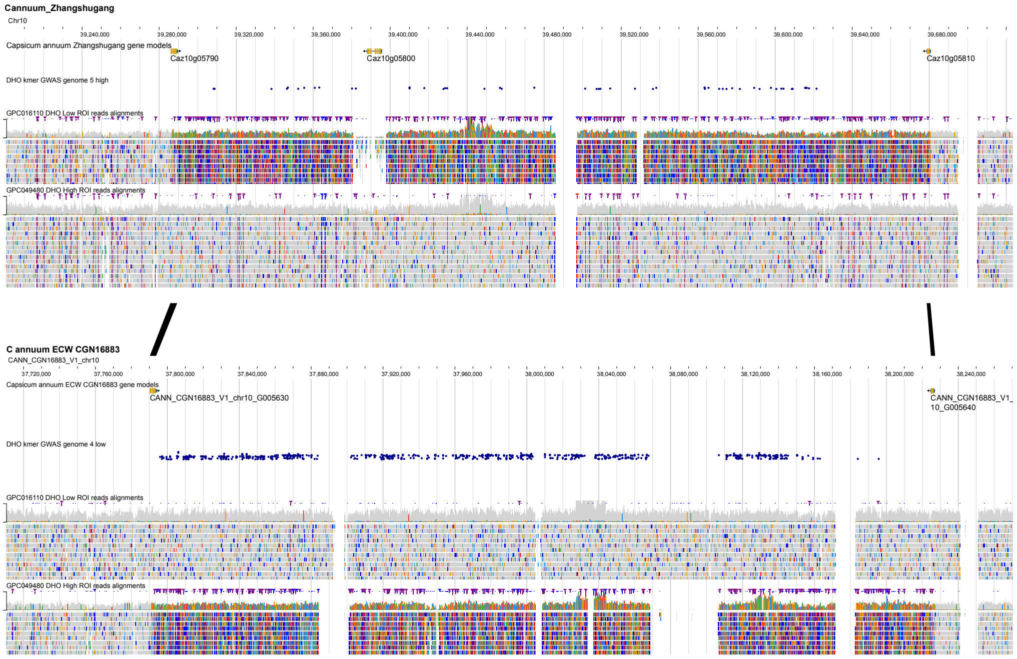


Figure S9: **Jbrowse2 overview of the dehydroxilinalool oxide significant locus** JBrowse 2 linear synteny view of the dehydroxilinalool oxide (DHO) significant locus regions in the *C. annuum* 'Zhangshugang' (top) and 'ECW' (bottom) genomes. Black bands connect genes assigned to the same homology groups. ROI-selected short reads are shown for accessions GPC049480 (high metabolite production), and GPC016110 (low metabolite production). Grey blocks and shading represent aligned reads and their coverage, while colored marks indicate sequence differences relative to the corresponding genome.

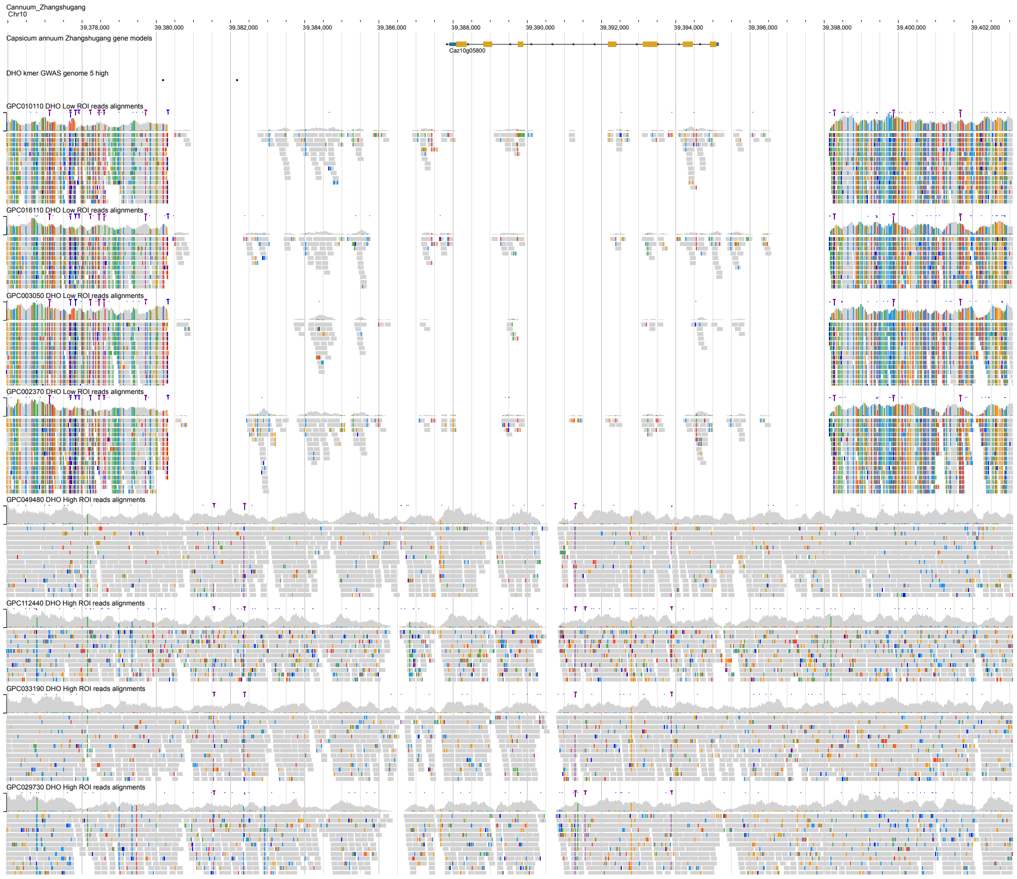


Figure S10: **Jbrowse2 overview of the variation of gene *Caz10g05800* in the dehydroxilinalool oxide significant locus** Jbrowse2 overview of read alignments of four accessions with high dehydroxilinalool oxide production (GPC049480, GPC0112440, GPC033190, GPC029730) and four accessions low dehydroxilinalool oxide production (GPC010110, GPC016110, GPC03050, GPC002370) against the *C. annuum* 'Zhangshugang' genome, including the significant $k$-mers located in the region.

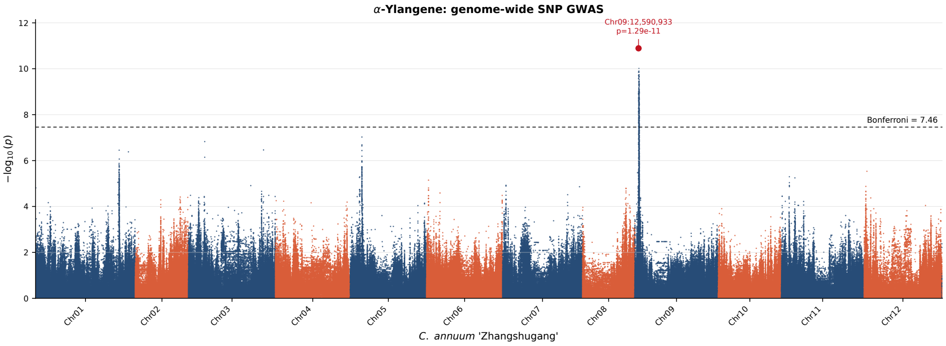


Figure S11: **Genome-wide SNP association results for $\alpha$-ylangene.** Manhattan plot of SNP genome-wide associations (GWAS) across the *C. annuum* 'Zhangshugang' genome. The dashed line indicates the Bonferroni-corrected genome-wide significance threshold ($-\log_{10}(p) = 7.46$), calculated from the number of SNP markers. The SNP highlighted in red on chromosome 2 represents the marker with the strongest statistical association.

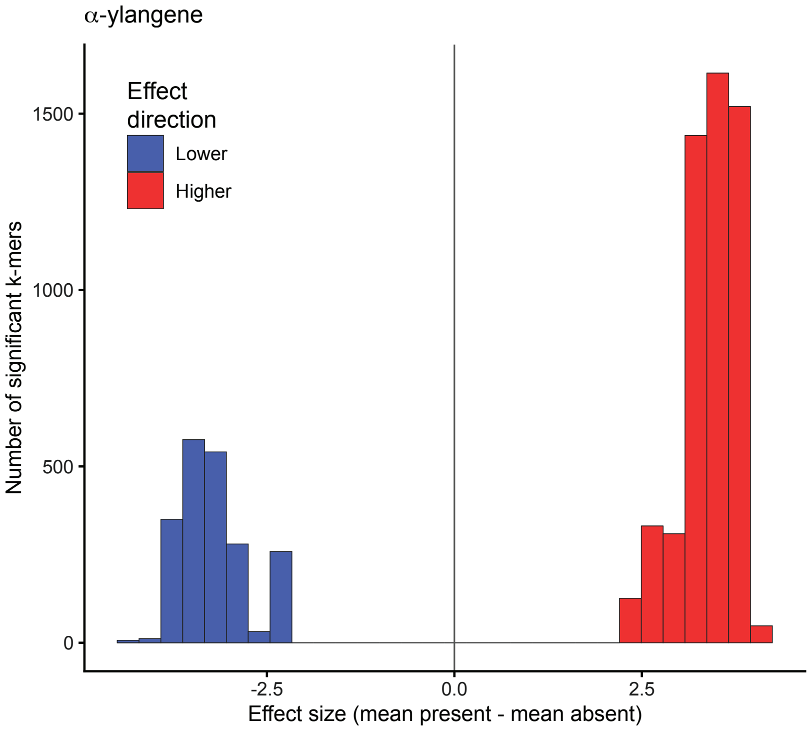


Figure S12: **Effect-size distribution of significant *k*-mers associated with $\alpha$-ylangene.** Histogram showing the difference in mean phenotype between accessions in which each significant *k*-mer was present and absent. Positive values indicate association with higher phenotype values, whereas negative values indicate association with lower phenotype values.

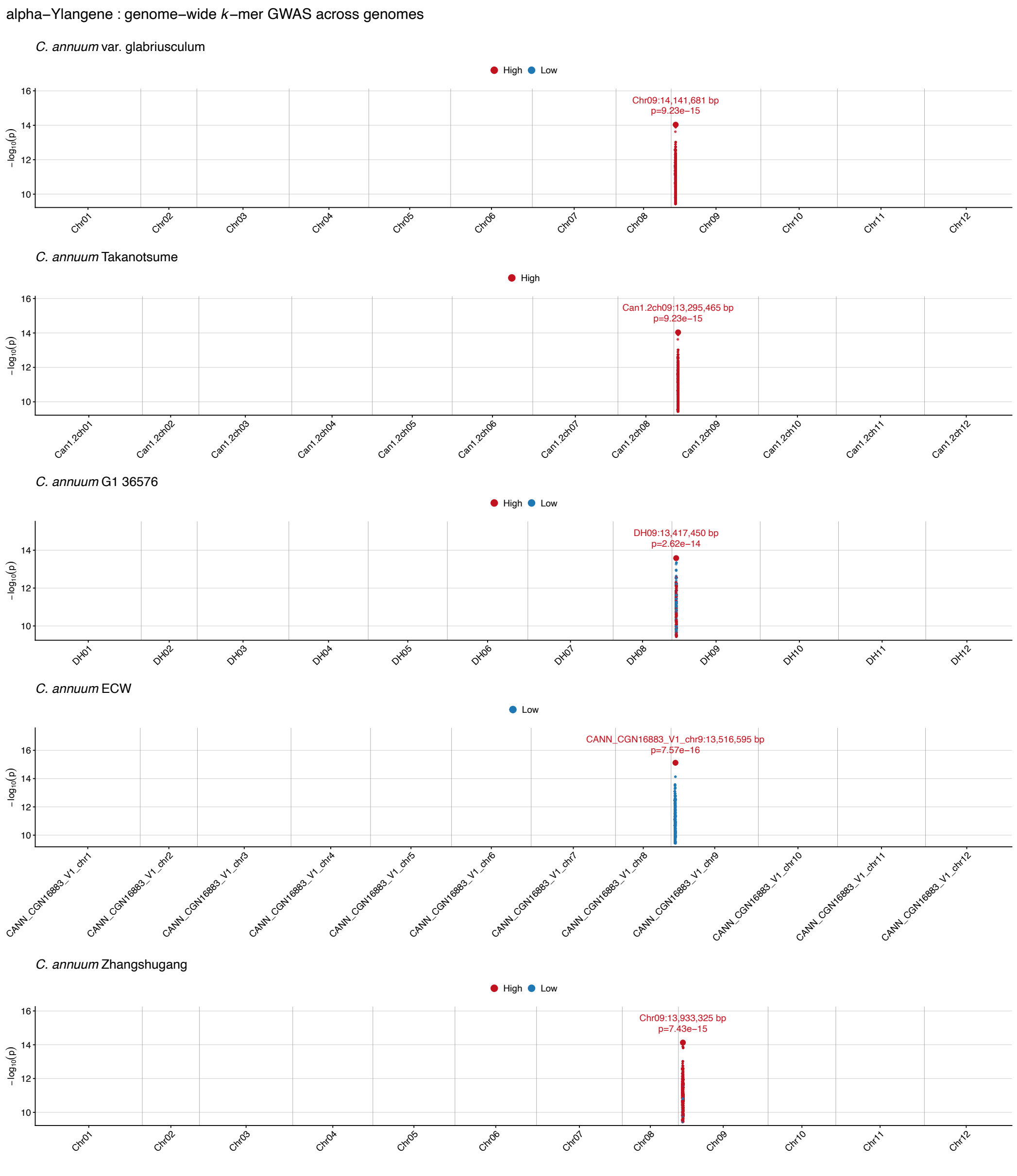


Figure S13: **Genome-wide $k$-mer association results for $\alpha$-ylangene.** Manhattan plots of $k$-mer genome-wide associations (GWAS) across the *C. annuum* genomes. Only significant $k$-mers are shown. The red highlighted $k$-mer marks the most significant association.

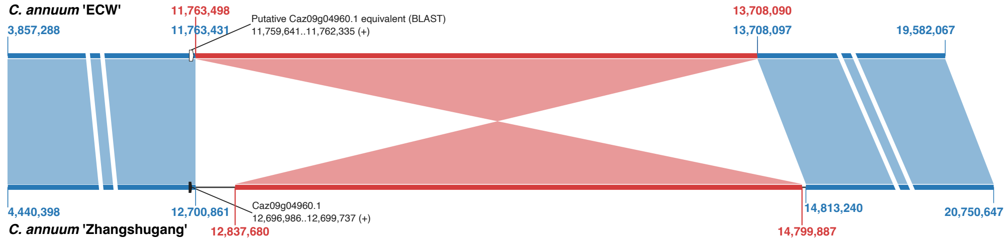


Figure S14: **Schematic representation of an inversion in the $\alpha$-ylangene locus between the *C. annuum* 'ECW' and 'Zhangshugang' genomes.** Blue regions are collinear between the two genomes, and red regions are inverted. Coordinates indicate the boundaries of the collinear and inverted alignment blocks. The terpene synthase gene *Caz09g04960*, located near the left inversion boundary in 'Zhangshugang', is shown. BLASTN of its genomic sequence (Chr09:12,696,986–12,699,737) against the 'ECW' genome identified a match at Chr09:11,759,641–11,762,335 (score = 4229.28; $E$ = 0; 93.8% identity; 2.6% gaps), supporting this unannotated region as the likely syntenic counterpart of *Caz09g04960*.

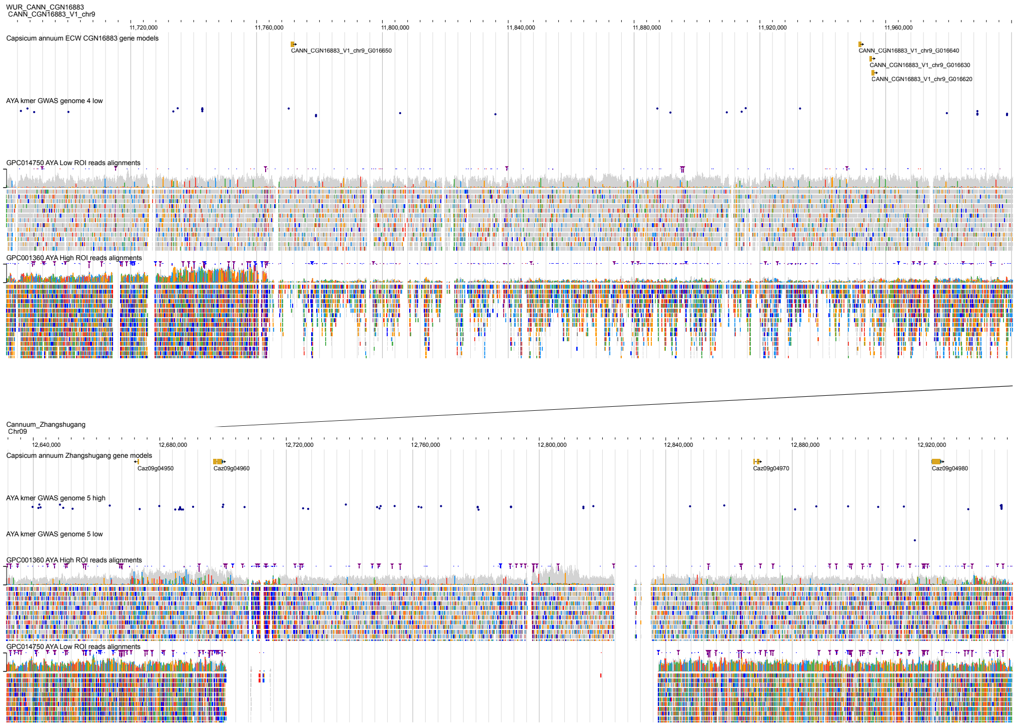


Figure S15: **Structural variation downstream of *Caz09g04960* in the $\alpha$-ylangene-associated locus.** JBrowse2 view of the left part of the associated locus in the *C. annuum* 'ECW' (top) and 'Zhangshugang' (bottom) genomes. The view includes *Caz09g04960*, the large deletion downstream of this gene, and the left breakpoint between the collinear and inverted regions. The region in 'ECW' with the highest sequence similarity to *Caz09g04960*, identified by BLAST, is shown in the corresponding upper track. Significant high- and low-associated *k*-mers and ROI-selected read alignments from GPC014750 (low) and GPC001360 (high) are shown.
